# Sexual and individual signatures are encoded in the temporal rate of Cape gannet display calls

**DOI:** 10.1101/2021.12.23.473090

**Authors:** Kezia Bowmaker-Falconer, Andrea Thiebault, Maёlle Connan, Thierry Aubin, Isabelle Charrier, Pierre Pistorius

**Author notes:** These authors contributed equally to this work. Kezia Bowmaker-Falconer / Andrea Thiebault.

## Abstract

Vocalisations play a vital role in animal communication, as they are involved in many biological functions. Seabirds often breed in large and dense colonies, making successful recognition between mates or between parents -and offspring crucial for reproductive success. Most seabird species, including Cape gannets (*Morus capensis*), are monomorphic and likely rely on acoustic signals for mate selection and mate recognition. This study aimed to better understand the use of vocalisations for sex and individual recognition in Cape gannets by describing the acoustic structure of their display calls at the nest. Vocalisations of nesting Cape gannets were recorded and acoustic measurements were extracted in both temporal and frequency domains. Values of the fundamental frequency and the average of Inter-Onset-Interval appeared to be the most important acoustic variables for sex determination. Both temporal and frequency parameters showed a potential for individual identity coding, with the average units’ Inter-Onset-Interval being the most important variable for individual identification for both sexes. This study provides the first evidence of sex-specific and individual vocal signatures in adult breeding Cape gannets. From an applied perspective, identified sex specific differences could potentially be used as a non-invasive method for field-based sex-determination in research and monitoring projects on Cape gannets.

## 2. Introduction

Communication facilitates and may even be vital to biological functions such as recognition [1], reproduction, foraging and defence [2]. Sex and individual identification can be essential for successful reproduction and can thus play an important role in an individual’s fitness. Differences in the vocalisations among individuals have been identified in both mammals [3,4] and birds [e.g. 5–9]. Vocal differences between sexes have also been detected in mammals [e.g. 10–12], anurans [e.g. 13,14] and birds [e.g. 6,7,9,15]. Studies on acoustic communication in birds have largely focused on terrestrial species with less but significant research focused on individual recognition in seabirds [8,15–21]. Different communication strategies thus exist in seabirds, each of which could be related to a specific breeding context [17].

Colonial animals, such as many seabirds, have developed specific acoustic recognition processes that assist with mate location and identification in particularly noisy and chaotic environments [15,22]. As central place foragers during the breeding season, seabirds alternate nest duties with foraging bouts at sea [23]. Identification of their partners and offspring on return to the colony is critical for successful reproduction [24]. Many seabirds are sexually monomorphic, suggesting that mechanisms other than visual cues are used for mate identification [25]. Indeed, vocal signals contain sexual and individual signatures in a number of seabird species, as shown in the Spheniscidae [e.g. 17,19], Laridae [e.g. 15,26,27], Procellariidae [e.g. 28,29] and Sulidae [e.g. 18,30] families. Often both temporal and frequency parameters play a role in the discrimination between sexes, as shown in black-legged-kittiwakes *Rissa tridactyla* [27], Yelkouan shearwaters *Puffinus yelkouan* [8,29] and blue-footed boobies *Sula nebouxii* [18]. For the display call of king penguins *Aptenodytes patagonicus*, the syntax of syllables is sex-specific and allows for a 100 % accuracy in sex determination [18].

Determining the sex of monomorphic seabirds in the field is often a challenge. Although this can potentially be achieved through observations during periods of copulation or when sex specific behaviours are undertaken (e.g. biting behaviour in male gannets [31]), the sex of study birds is often established through laboratory-based molecular work [32,33]. This has the disadvantage of being invasive as samples (usually blood or feathers for birds) need to be collected from captured individuals. The samples can then only be processed afterwards, which can be problematic when individuals of a particular sex need to be targeted (e.g. for tracking studies). This technique is also costly as samples need to be analysed in a laboratory by trained professionals [7,34]. Therefore, the use of a more time-efficient and non-invasive technique for sexing seabirds in the field is desired, such as through their call characteristics [19,29].

Within the family Sulidae, individual signatures in the vocalisations of Northern *Morus bassanus* and Australasian *Morus serrator* gannets have been documented [30,35]. Sexspecific differences, on the other hand, remain to be studied in Northern gannets, yet seems to be absent in Australasian gannets [30]. The potential for either individual or sex-specific signatures in the third gannet species, the Cape gannets are yet to be investigated. The Cape gannet is an endangered [36] species endemic to southern Africa, which like other members of the Sulidae family congregates in large, dense colonies during the breeding season [37]. Over the last 20 years, the Cape gannet population has declined by 52 % across its six breeding colonies in South Africa [38]. They are largely sexually monomorphic despite slight differences in gular stripe length, which cannot be used for reliable sex identification, allowing only 65 % of correct classifications [39]. Acoustic analysis of the vocalisation emitted at the nest can thus potentially help determine if individuals and sex can be identified in the field, making research, which informs conservation management, increasingly effective.

At their breeding colony, Cape gannets produce a number of vocalisations in specific behavioural contexts (e.g. when landing, meeting with their partner, leaving the nest and fighting [31]). In this study, we focused on the display vocalisations, potentially important for partner recognition [31]. The mutual display (or ‘Mutual Greeting’ as per [31]) is a ceremony during which the two partners face each other in a synchronised dance with associated vocalisations [31]. This dance is thus performed as a duet, not only during courtship or mating, but also every time they meet again on the nest during the breeding season, suggesting an important role for sexual and individual recognition. However, during the mutual display, the calls of each partner overlap, preventing an accurate acoustic analysis. The same behaviour is also performed solitarily, putatively as a form of territorial behaviour or a nest ownership display (the ‘Solo Bow’ as per [31]). For this study we analysed the single display calls, produced during the ‘Solo Bow’ behaviour.

This study aims to better understand the potential use of vocalisations for sex and individual recognition in Cape gannets by 1) describing the acoustic characteristics of the single display call (henceforth referred to as display call) of nesting Cape gannets, 2) determining if there are sex-specific vocal features in these calls, potentially allowing for field-based sex determination and, 3) assessing the occurrence of individual vocal signatures in the display calls.

## 3. Materials and Methods

### Data collection

Data were collected on Cape gannets during their brooding phase in December 2015 on Bird Island (33°50□26□S 26°17□10□E, Algoa Bay, South Africa), which holds the largest breeding colony of Cape gannets with more than 90 000 breeding pairs [37]. Two clumps of twenty Cape gannet nests each were marked with unique numbers and these were mapped. At least one partner per nest was marked using short-term animal marking sticks for individual identification. In addition for some of these nests, a breeding adult was captured using a pole with a hook as part of another study [20] and a couple of breast feathers were plucked for sex identification based on DNA analyses.

Over a two-week period, the vocalisations and associated behaviour of breeding adults from the study nests were recorded daily for approximately 2-3 h during the early morning or late afternoon, when gannet nest activities are typically relatively high [40]. Vocalisations were recorded using a microphone Beyer-Dynamic M 69 TG (frequency response: 50 Hz-16 kHz ± 2.5 dB) connected to a digital recorder Zoom H4N (sampling frequency 44.1 kHz). The microphone was placed ~1 m from the study nests for recording purposes. A long cable allowed the observer to lie at ~5 m distance from the colony, thus minimizing potential observer effects. The identity of vocalising birds, together with their behaviour when vocalising were manually noted by a single observer throughout fieldwork (AT). Annotated behaviours with associated vocalisations included: landing and returning to nest, leaving the nest, mutual display (or ‘Mutual Greeting’ as per [31]), single display (‘Solo bow’ as per [31]) and fighting (two gannets grabbing each other’s beaks). In addition, a video camera recorded the monitored nests to allow for further behavioural observations during data analyses.

### Molecular sexing

Genomic DNA was extracted from the plucked feathers using a Chelex extraction method, implemented previously on Cape gannets [39]. Fragments of the sex-linked CHF-1 gene were amplified using 2550F (5’-GTTACTGATTC GTCTACGAGA-3’) and 2718R (5’ - TTGAAATGATCCAGTGCTTG-3’) primers, with females revealing in agarose gel as two fragments (ZW) and males as a single fragment (ZZ) [41]. Polymerase chain reactions in a 15 μL solution containing 7.5 μL GoTaq^®^ G2 Hot Start Green Master Mix (Promega), 0.4 μmol of each primer and 46 – 247 ng of genomic DNA were performed using a C1000 Touch Thermal Cylinder (BioRad). Initial denaturing of DNA was set at 94°C for 2 min, followed by 41 cycles with a denaturation step at 94°C for 30 s, an annealing step at 50°C for 30 s and an extension step at 72°C for 45 s. A final extension step of 5 min at 72°C was added after the last cycle. PCR products (5 μL) were separated on a 1.8% agarose gel with 1X TAE buffer. After electrophoresis at 100 V for 30 min, gels were stained with GelRed™ Nucleic Acid Gel stain (Biotium) and bands were visualized under ultraviolet light.

### Measure of acoustic variables

To increase the precision of frequency measurements, all the calls were resampled at 16 kHz and analysed using Avisoft-SASLab Pro (version 5.2, Avisoft Bioacoustics, Germany). A call was defined as temporally distinct sounds associated with a display dance behaviour (Fig. 1, Supplementary Video 1). A call was divided into up-movement and down-movement parts according to the head movement of the bird (successively facing up and down, as observed from the recorded videos), with each up-movement and down-movement parts composed of several sound units (Fig. 1, Supplementary Video 1). Calls were selected for measurements wherever the quality of recordings allowed (i.e. low background noise and no overlap with other calls). Among the monitored and recorded birds, we selected for the acoustic analysis only the individuals for which a minimum of four calls were recorded with sufficient acoustic quality (number four arbitrarily chosen as a trade-off between a reasonable number of calls per individual and a reasonable number of individuals kept for the analyses). For each selected bird, a maximum of six calls per individual (selected randomly) were kept for acoustic measurements to limit imbalance in the dataset. Acoustic variables were measured on one of each up-movement and down-movement parts, selected in the middle of the entire call to ensure full momentum of the behaviour, as well as on the first unit of each measured part. A total of 36 acoustic variables were measured (12 temporal and 24 frequency features). From the oscillogram, the duration (in s) of the part and of the first unit of each part were measured, as well as the number of units in each part (separated by ~0.1 s strong amplitude declines) and the number of segments in a unit (separated by ~0.01 s by strong amplitude declines; Fig. 1). From the average energy spectrum displayed between 0.3 and 5 kHz, the fundamental frequency (F0, Hz), the frequency of maximum amplitude (Fmax, Hz), the first (Q25, Hz), the second (Q50, Hz) and the third (Q75, Hz) quartiles were measured automatically, as well as the percentage of energy occurring below 1200 Hz (E1200). In addition, for units the pulse rate (Hz) of sound units was automatically extracted (using the ‘Pulse train analyses’ function in Avisoft-SASLab Pro) as a measure of the temporal rate of segment emission within a unit. The temporal rate of sound production was also evaluated over an entire up-and-down sequence using the measure of Inter-Onset-Intervals (IOI), defined as the “time elapsed between the beginning of one event (i.e. onset) and the beginning of the next event” [42]. The average and standard deviation of IOI measured between successive units were calculated for each measured call.

**Figure 1:**
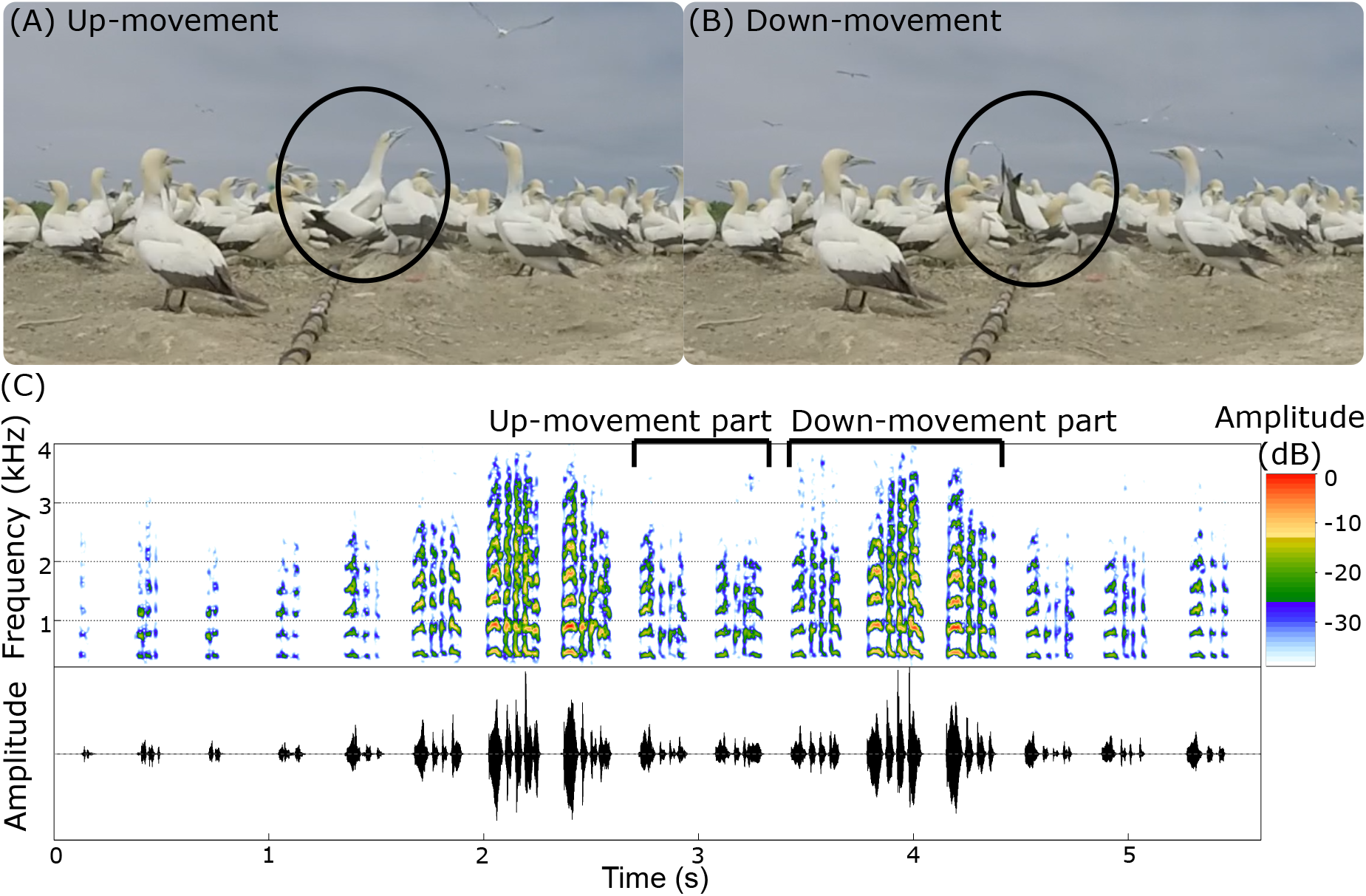
Illustration of a display call produced by a Cape gannet breeding on Bird Island (Algoa Bay, South Africa). Characteristic up (A) and down (B) movement associated with the call. Images extracted from videos. (C) Representation of the sound produced during an entire display call, comprising of successive up-movement and down-movement parts, including the spectrogram, i.e. frequency over time (top row) and the oscillogram, i.e. amplitude over time (bottom row). Figure generated using the ‘Seawave’ package [62] in R software.

### Sexual dimorphism in display calls

The mean, standard deviation and range of all 36 acoustic variables were calculated and compared between sexes. The distribution of each acoustic variable was tested for normality using a Shapiro-Wilk test. As the majority of them were not normally distributed, the distribution between sexes were compared using non-parametric tests. The variances of distributions were compared using a Fligner-Killeen test of homogeneity of variance and their medians using a Kruskal-Wallis rank sum test. The acoustic structure of calls emitted by the two sexes was then compared in a multivariate analysis. Only variables for which at least one of the Kruskal-Wallis or Fligner-Killeen statistical tests resulted in significant differences were kept (21 acoustic variables). The random forest algorithm (RF) was chosen because it does not require assumptions on the distribution of predictor variables [43]. The global accuracy of prediction is estimated intrinsically in the algorithm using a bootstrap process and calculated as a proportion of correct classification. In addition, we used the indicator “precision” [44] to calculate the number of correct predictions per class (sexes), based on the confusion matrix. We then compared this accuracy of prediction per class to a prediction by chance, calculated as the number of calls in the class (male or female) divided by the total number of calls (following the method in [20]). This allows us to evaluate the strength of the prediction in comparison to a random allocation of class based on occurrences. Furthermore, the bootstrap process in the RF algorithm can be used to estimate the importance of variables for predictions. This was used to identify the acoustic variables contributing the most to the sex identification. Collinearity between variables was tested since a high collinearity between two important variables may affect their ranking in the list of important variables. Three couples of variables were found with a high collinearity (>0.9). For each couple, one variable was removed (the most difficult one to interpret). This resulted in a set of 18 acoustic variables included in the RF to compare sexes.

The RF was run in R software using the package “randomForest” [45]. The number of trees to be grown from bootstrap samples of the dataset (parameter “ntree”) was set at 200. This ensured convergence of the results (Supplementary Fig. 1) as well as robustness in the measure of variable importance [46]. To set the number of variables to be randomly selected at each node (parameter “mtry”) we used the default value for classification: the square root of the total number of variables (18), so four in our case.

### Individual signatures in display calls

The individual signatures were studied within sexes, allocating a sex to birds where no samples were collected for molecular-sexing, based on the acoustics of their display calls (see results on sexual dimorphism). For each sex, we assessed the potential of individual coding (PIC) for each of the 36 acoustic variables by dividing the coefficient of variation between individuals (CV inter-individual) calculated on all individuals pooled together with the mean of CVs calculated for each individual (CV intra and inter-individual) [47]. The CV was calculated according to the formula for small samples sizes: CV={100(SD/Xmean)[1+(1/4n)]}, where SD is the standard deviation, Xmean the mean for each individual and n the number of calls per individual [48]. A PIC value greater than 1 means that the inter-individual variability is greater than the intra-individual variability and so the given variable can be interpreted as individual-specific. In addition, the distribution of each variable per individual was compared using a Kruskal-Wallis rank sum test.

Individual identity can be coded from a combination of variables, so the set of acoustic variables was then compared per individual using a multivariate analysis. The RF algorithm was used to classify the acoustic structure of calls per individual following the same method as explained in the section on sexual dimorphism, but different sets of variables were included in the different models, depending on the univariate statistical results and the collinearity between variables. For males, all variables were considered since they all resulted in a significant difference according to the Kruskal-Wallis test. Among these, five couples of variables were found to be highly correlated (>0.9) so five variables were removed from the set to reduce collinearity and improve the identification of important variables. This resulted in a total of 31 acoustic variables included in the RF for males. Consequently, the algorithm for individual differences among males was applied with the parameter “mtry” set at six (square root of 31) and with the parameter “ntree” set at 4000 to ensure convergence of the results (Supplementary Fig. 1). For females, correlation was tested among the 18 variables that showed significant differences among individuals according to the Kruskal-Wallis test. Only one couple of variables was highly correlated (>0.9) resulting in a total of 17 acoustic variables included in the RF comparing individuals among females. Consequently, the RF algorithm for females was applied with the parameter “mtry” set at four (square root of 17) and with the parameter “ntree” set at 1000 to ensure convergence of the results (Supplementary Fig. 1).

## 4. Results

A total of 184 display calls was recorded, among which 97 were produced by molecular-sexed gannets (74 calls from six males and 23 calls from four females). From these recordings, acoustic measurements were taken from a total number of 80 calls, which were comprised of sixteen different individuals including six males (4-6 calls per individual totalling 31 calls), four females (4-6 calls per individual totalling 19 calls) and six unsexed individuals (five calls per individual totalling 30 calls).

### The display call of Cape gannets

The display call was always associated with a characteristic up (A) and down (B) movement (Fig. 1, Supplementary Video 1). It was composed of a series of distinct sound units (separated by strong amplitude declines of ~0.1 s) emitted successively throughout the dance (Fig. 1c). The up and down movement was typically repeated two or three times (up to four times) during the whole display behaviour. Each up and each down part were composed of a specific number of sound units (ranging between two and eight), and each unit was furthermore composed of a series of segments, separated by ~0.01 s strong amplitude declines. The number of sound units within each part, as well as the number of segments within units (ranging between two and seven), varied among individuals. Fig. 1c shows the spectrogram of the up-movement part of a given individual composed of two units, whilst the down-movement part was comprised of three units. Within the units of the up-movement part the first unit was comprised of four segments and within the units of the down-movement part the first unit was also comprised of four segments.

### Sexual dimorphism in display calls

To assess acoustic variations between sexes in the display calls of Cape gannets, we analysed 31 calls from molecular-sexed males and 19 calls from molecular-sexed females (4-6 calls per individual). The variables showing the highest statistical differences between sexes (p<0.001 for both tests on median and variance) were the IOIm, the fundamental frequency (measured on the up-movement part and on the first unit of the down-movement part), and the duration of the first unit of both up-movement and down-movement parts (Table 1). Overall, variables showed more differences in terms of the median of distribution than the variance, with 20 and 10 variables significantly different according to the Kruskal-Wallis and the Fligner-Killeen tests, respectively (Table 1). The majority (7/12) of acoustic variables related to the temporal domain (e.g. IOIm, number and duration of units, number of segments, pulse rate) showed high significant differences (p<0.01), whereas for frequency parameters, only the fundamental frequency showed a high significant difference between sexes (p<0.001, Table 1). Interestingly, even if the IOIm (units’ temporal rate) was different between males and females, with a faster tempo in females than in males (Supplementary Audios 1-4), the unit rate seemed consistent for both sexes as shown by the small value of IOIsd.

**Table 1.**
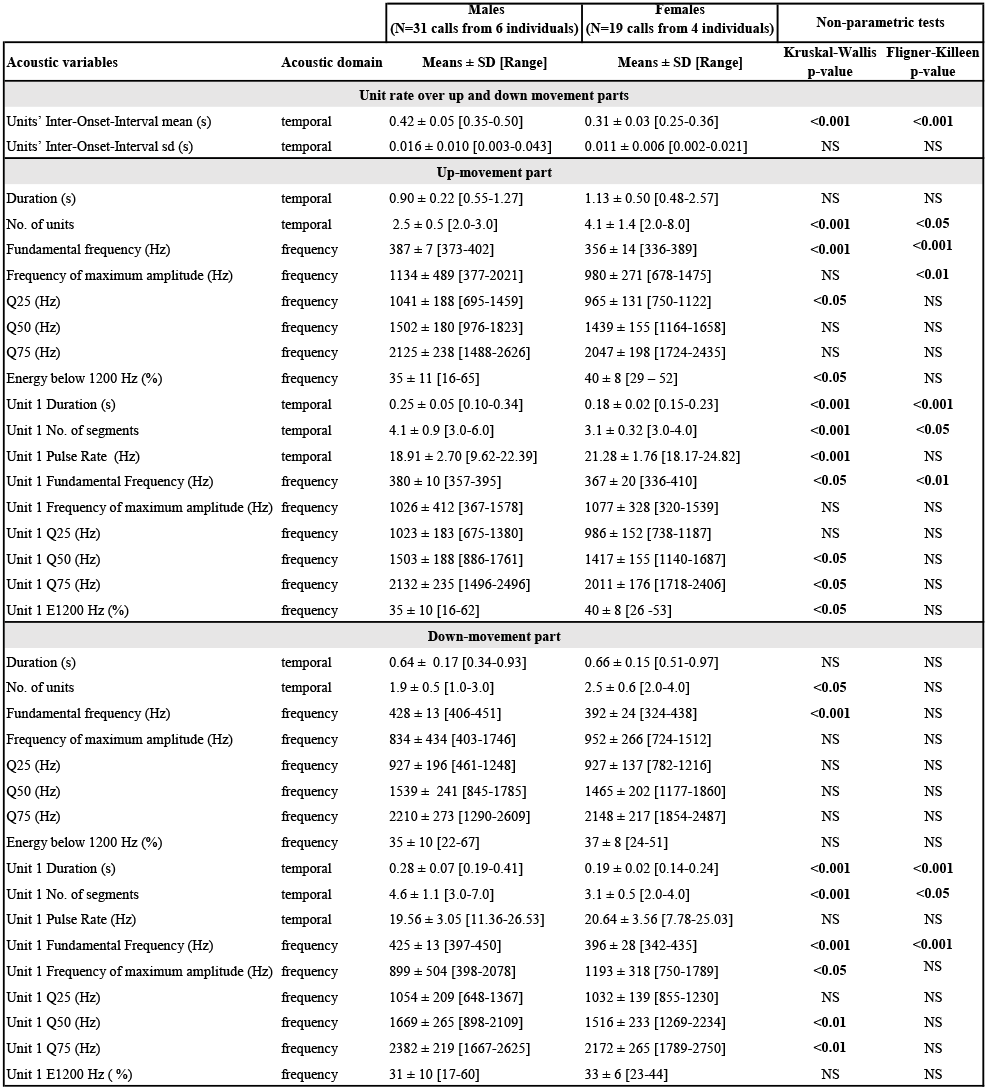
Summary of distribution of acoustic variables measured on the display calls produced by breeding male and female Cape gannets. Differences in variance of distribution per context were evaluated using the Fligner–Killeen test of homogeneity of variance. Differences in median of distribution per variable were evaluated using the Kruskal–Wallis rank sum test. Q25, Q50, Q75 = first, second and third quartile of energy distribution. E1200 = percentage of energy occurring below 1200 Hz.

|  |  | Males<br>(N=31 calls from 6 individuals) | Females<br>(N=19 calls from 4 individuals) | Non-parametric tests |  |
| --- | --- | --- | --- | --- | --- |
| Acoustic variables | Acoustic domain | Means ± SD [Range] | Means ± SD [Range] | Kruskal-Wallis<br>p-value | Fligner-Killeen<br>p-value |
| Unit rate over up and down movement parts |  |  |  |  |  |
| Units' Inter-Onset-Interval mean (s) | temporal | 0.42 ± 0.05 [0.35-0.50] | 0.31 ± 0.03 [0.25-0.36] | <0.001 | <0.001 |
| Units' Inter-Onset-Interval sd (s) | temporal | 0.016 ± 0.010 [0.003-0.043] | 0.011 ± 0.006 [0.002-0.021] | NS | NS |
| Up-movement part |  |  |  |  |  |
| Duration (s) | temporal | 0.90 ± 0.22 [0.55-1.27] | 1.13 ± 0.50 [0.48-2.57] | NS | NS |
| No. of units | temporal | 2.5 ± 0.5 [2.0-3.0] | 4.1 ± 1.4 [2.0-8.0] | <0.001 | <0.05 |
| Fundamental frequency (Hz) | frequency | 387 ± 7 [373-402] | 356 ± 14 [336-389] | <0.001 | <0.001 |
| Frequency of maximum amplitude (Hz) | frequency | 1134 ± 489 [377-2021] | 980 ± 271 [678-1475] | NS | <0.01 |
| Q25 (Hz) | frequency | 1041 ± 188 [695-1459] | 965 ± 131 [750-1122] | <0.05 | NS |
| Q50 (Hz) | frequency | 1502 ± 180 [976-1823] | 1439 ± 155 [1164-1658] | NS | NS |
| Q75 (Hz) | frequency | 2125 ± 238 [1488-2626] | 2047 ± 198 [1724-2435] | NS | NS |
| Energy below 1200 Hz (%) | frequency | 35 ± 11 [16-65] | 40 ± 8 [29 – 52] | <0.05 | NS |
| Unit 1 Duration (s) | temporal | 0.25 ± 0.05 [0.10-0.34] | 0.18 ± 0.02 [0.15-0.23] | <0.001 | <0.001 |
| Unit 1 No. of segments | temporal | 4.1 ± 0.9 [3.0-6.0] | 3.1 ± 0.32 [3.0-4.0] | <0.001 | <0.05 |
| Unit 1 Pulse Rate (Hz) | temporal | 18.91 ± 2.70 [9.62-22.39] | 21.28 ± 1.76 [18.17-24.82] | <0.001 | NS |
| Unit 1 Fundamental Frequency (Hz) | frequency | 380 ± 10 [357-395] | 367 ± 20 [336-410] | <0.05 | <0.01 |
| Unit 1 Frequency of maximum amplitude (Hz) | frequency | 1026 ± 412 [367-1578] | 1077 ± 328 [320-1539] | NS | NS |
| Unit 1 Q25 (Hz) | frequency | 1023 ± 183 [675-1380] | 986 ± 152 [738-1187] | NS | NS |
| Unit 1 Q50 (Hz) | frequency | 1503 ± 188 [886-1761] | 1417 ± 155 [1140-1687] | <0.05 | NS |
| Unit 1 Q75 (Hz) | frequency | 2132 ± 235 [1496-2496] | 2011 ± 176 [1718-2406] | <0.05 | NS |
| Unit 1 E1200 Hz (%) | frequency | 35 ± 10 [16-62] | 40 ± 8 [26 -53] | <0.05 | NS |
| Down-movement part |  |  |  |  |  |
| Duration (s) | temporal | 0.64 ± 0.17 [0.34-0.93] | 0.66 ± 0.15 [0.51-0.97] | NS | NS |
| No. of units | temporal | 1.9 ± 0.5 [1.0-3.0] | 2.5 ± 0.6 [2.0-4.0] | <0.05 | NS |
| Fundamental frequency (Hz) | frequency | 428 ± 13 [406-451] | 392 ± 24 [324-438] | <0.001 | NS |
| Frequency of maximum amplitude (Hz) | frequency | 834 ± 434 [403-1746] | 952 ± 266 [724-1512] | NS | NS |
| Q25 (Hz) | frequency | 927 ± 196 [461-1248] | 927 ± 137 [782-1216] | NS | NS |
| Q50 (Hz) | frequency | 1539 ± 241 [845-1785] | 1465 ± 202 [1177-1860] | NS | NS |
| Q75 (Hz) | frequency | 2210 ± 273 [1290-2609] | 2148 ± 217 [1854-2487] | NS | NS |
| Energy below 1200 Hz (%) | frequency | 35 ± 10 [22-67] | 37 ± 8 [24-51] | NS | NS |
| Unit 1 Duration (s) | temporal | 0.28 ± 0.07 [0.19-0.41] | 0.19 ± 0.02 [0.14-0.24] | <0.001 | <0.001 |
| Unit 1 No. of segments | temporal | 4.6 ± 1.1 [3.0-7.0] | 3.1 ± 0.5 [2.0-4.0] | <0.001 | <0.05 |
| Unit 1 Pulse Rate (Hz) | temporal | 19.56 ± 3.05 [11.36-26.53] | 20.64 ± 3.56 [7.78-25.03] | NS | NS |
| Unit 1 Fundamental Frequency (Hz) | frequency | 425 ± 13 [397-450] | 396 ± 28 [342-435] | <0.001 | <0.001 |
| Unit 1 Frequency of maximum amplitude (Hz) | frequency | 899 ± 504 [398-2078] | 1193 ± 318 [750-1789] | <0.05 | NS |
| Unit 1 Q25 (Hz) | frequency | 1054 ± 209 [648-1367] | 1032 ± 139 [855-1230] | NS | NS |
| Unit 1 Q50 (Hz) | frequency | 1669 ± 265 [898-2109] | 1516 ± 233 [1269-2234] | <0.01 | NS |
| Unit 1 Q75 (Hz) | frequency | 2382 ± 219 [1667-2625] | 2172 ± 265 [1789-2750] | <0.01 | NS |
| Unit 1 E1200 Hz (%) | frequency | 31 ± 10 [17-60] | 33 ± 6 [23-44] | NS | NS |

The RF classification for the two sexes showed a global accuracy of prediction of 98% with a near perfect classification. The indicators precision showed that 95 % (18/19) of female calls and 100 % (31/31) of male calls were correctly classified. These predictions were 2.5 and 1.6 times better than a prediction by chance for females and males, respectively (Table 2).

**Table 2.**
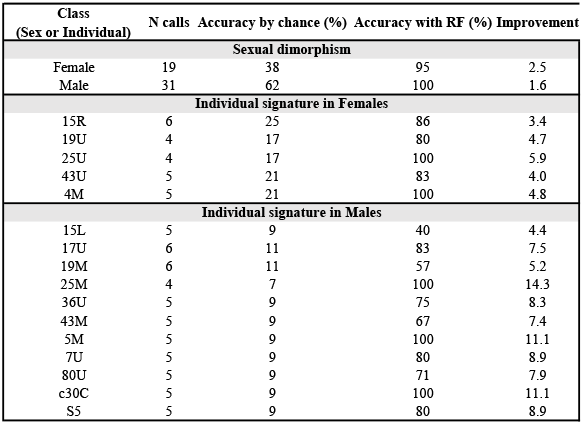
Comparison of the accuracy of prediction obtained by chance (number of calls in a class divided by the total number of calls in the given analysis) or prediction using the random forest algorithm (indicator precision), for the analysis on sexual dimorphism (classifying females or males) and for the analysis on individual signatures among females or males (classifying individuals).

The most important variables to correctly predict the sex of an individual from its display call was by far the IOIm (units’ temporal rate), with a mean decrease in accuracy of >10% when this variable was not included (Fig. 2). The second most important variable was the fundamental frequency during the up-movement part (Fig. 2). The following three important variables to correctly predict the sex of an individual still related to the fundamental frequency (during the down-movement part) and temporal variables measured on units (number of units and duration of the first unit in the up-movement part, Fig. 2). Among the 18 acoustic variables included in the RF comparing sexes, six out of the seven temporal variables appeared in the top ten most important variables. In comparison, only four out of the eleven frequency variables appeared in the top ten, with all four being measures of fundamental frequencies on different parts of the call (Fig. 2).

**Figure 2.**
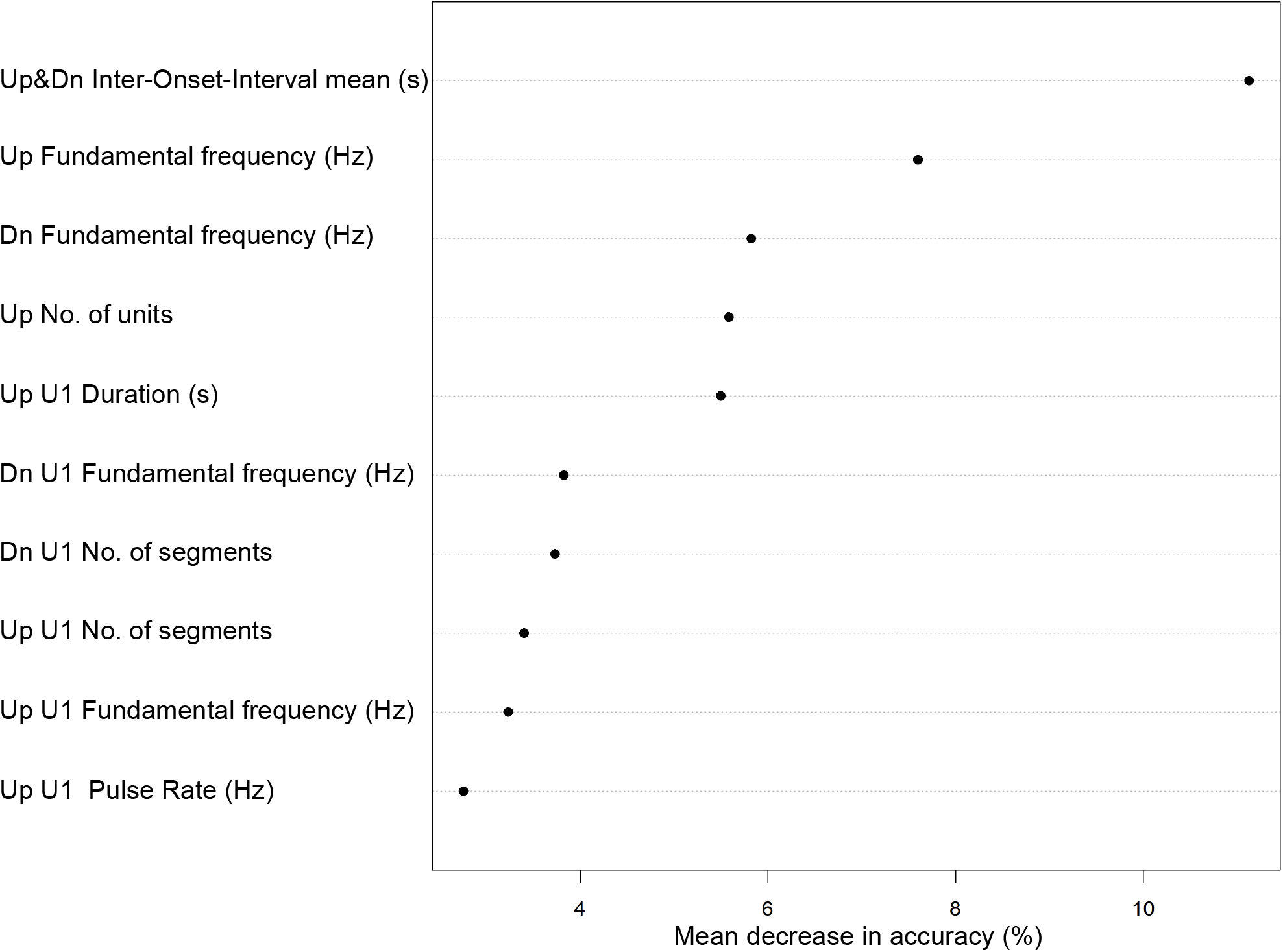
Analyses on sex-specific signatures in the Cape gannet display calls: ranking of the importance of acoustic variables for sex determination, calculated as a mean decrease in accuracy in the random forest algorithm. Only the first 10 variables are shown here. Up = up-movement part. Dn = down-movement part. U1 = first unit.

Compared to males, females had a lower fundamental frequency for both up-movement and down-movement parts (up-movement part average 356 Hz vs 387 Hz for females and males respectively, down-movement part average 392 Hz vs 428 Hz for females and males respectively, Table 1). Females produced a higher number of sound units (up-movement part average 4.1 vs 2.5 units for females and males, respectively, down-movement part average 2.5 vs 1.9 for females and males respectively, Table 1) at a higher temporal rate (IOI mean 0.31 vs 0.42 for females and males respectively, Table 1, Supplementary Audios 1-4).

Ultimately, the two most important variables identified by the RF algorithm, were IOIm and F0 during the up-movement part, which seemed sufficient to distinguish the sex of a Cape gannet based on its display call (Fig. 3). Two thresholds could be identified (380 Hz for the UpF0 and 0.35 s for the IOIm, Fig. 3) and if used simultaneously they allowed to successfully discriminate with 100 % accuracy the sex of the Cape gannet. Following this method, we were able to identify one female and five males among the sex-unknown recorded individuals.

**Figure 3.**
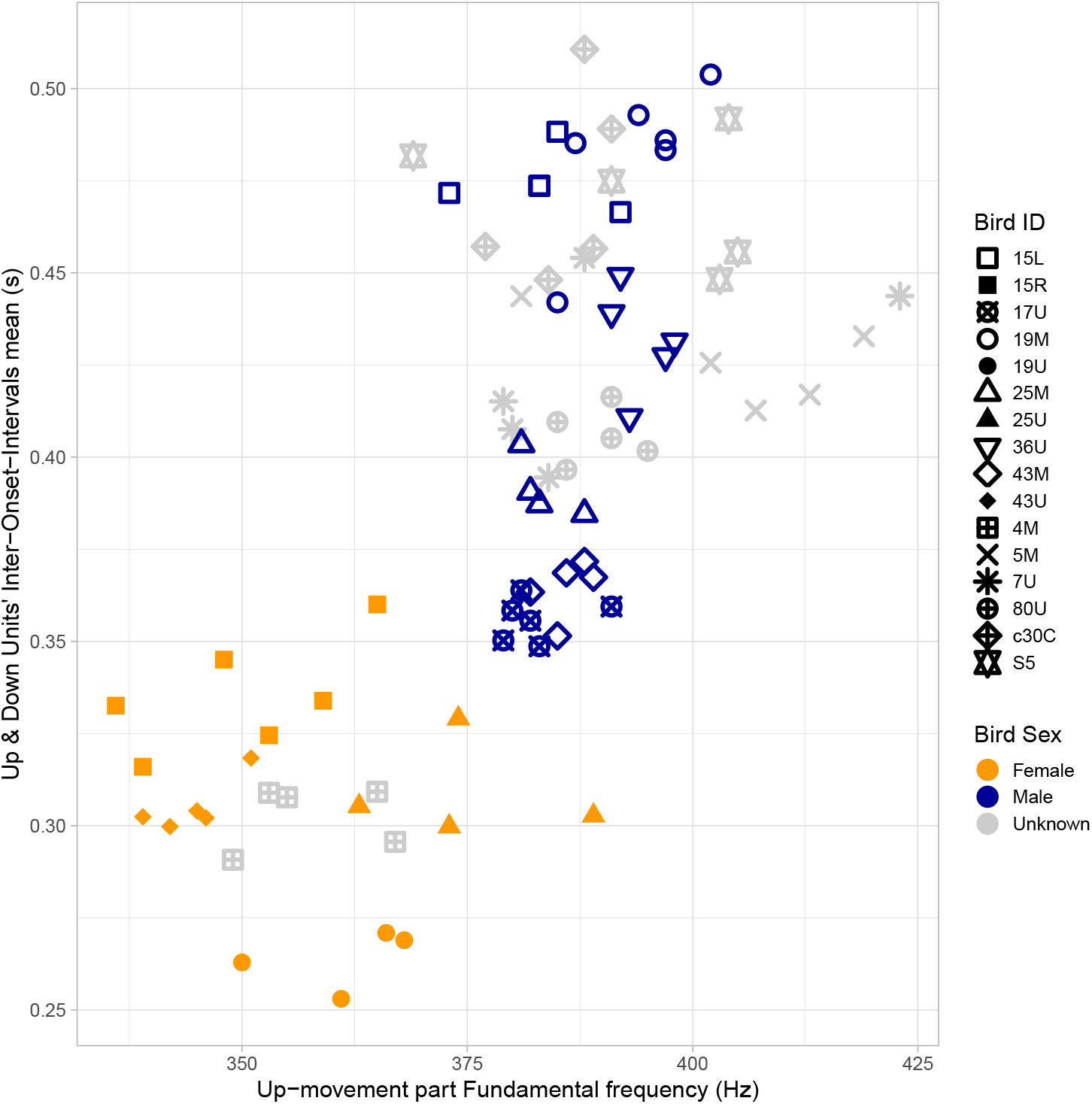
Mean Inter-Onset-Intervals measured along the up and down sequence, as a function of the fundamental frequency during the up-movement part in the display call of Cape gannets. Bird ID = individual identification of different birds. Females (filled orange symbols) and males (dark blue open symbols) were genetically sexed.

### Individual signature in display calls

The individual vocal signatures were assessed separately for each sex, using the entire data set which included five females (four molecular-sexed and one acoustically-sexed) and 11 males (six molecular-sexed and five acoustically-sexed). For males, all of the acoustic variables measured showed PIC values greater than one, with significant differences between individuals (Kruskal-Wallis test p<0.001, n = 56 calls from 11 individuals, Table 3). For females, the majority of the variables also showed PIC values greater than one, but not all of them (24 out of 36 variables with a PIC < 1.1), with 18 of them also showing significant differences (Kruskal-Wallis test, p<0.01 or p<0.05, n = 24 calls from five individuals). The variable with the highest PIC for both males and females was the IOIm, with a PIC value of 3.2 and 2.3 respectively. Other variables with PIC values greater than two included the number of units in the up-movement part and the duration of the first unit during the down-movement part, both for males only (Table 3).

**Table 3.** Analyses of the individual vocal signatures in the display call of males and females Cape gannets. Potential for individual coding (PIC) were calculated for each acoustic variable. The difference in median of distribution per variable evaluated using the Kruskal– Wallis rank sum test. NS = non-significant. A PIC value >1 indicates a potential for individual coding, with the highest the value is, the highest the potential is. Q25, Q50, Q75 = first, second and third quartile of energy distribution. E1200 = percentage of energy occurring below 1200 Hz.

|  |  | Males<br>(N=56 calls from 11 individuals) |  | Females<br>(N=24 calls from 5 individuals) |  |
| --- | --- | --- | --- | --- | --- |
| Acoustic variables | Acoustic domain | PIC | Kruskal-Wallis<br>p-value | PIC | Kruskal-Wallis<br>p-value |
| Unit rate over up and down movement parts |  |  |  |  |  |
| Units' Inter-Onset-Interval mean (s) | temporal | 3.22 | <0.001 | 2.30 | <0.01 |
| Units' Inter-Onset-Interval sd (s) | temporal | 1.35 | <0.001 | 1.04 | NS |
| Up-movement part |  |  |  |  |  |
| Duration (s) | temporal | 1.57 | <0.001 | 1.52 | <0.05 |
| No. of units | temporal | 2.03 | <0.001 | 1.38 | <0.05 |
| Fundamental frequency (Hz) | frequency | 1.32 | <0.001 | 1.48 | <0.01 |
| Frequency of maximum amplitude (Hz) | frequency | 1.13 | <0.001 | 1.49 | <0.05 |
| Q25 (Hz) | frequency | 1.32 | <0.001 | 1.06 | NS |
| Q50 (Hz) | frequency | 1.58 | <0.001 | 0.97 | NS |
| Q75 (Hz) | frequency | 1.44 | <0.001 | 1.06 | NS |
| Energy below 1200 Hz (%) | frequency | 1.59 | <0.001 | 1.04 | NS |
| Unit 1 Duration (s) | temporal | 1.87 | <0.001 | 1.58 | NS |
| Unit 1 No. of segments | temporal | 1.32 | <0.001 | 1.35 | NS |
| Unit 1 Pulse Rate (Hz) | temporal | 1.19 | <0.001 | 1.05 | NS |
| Unit 1 Fundamental Frequency (Hz) | frequency | 1.19 | <0.001 | 1.42 | <0.05 |
| Unit 1 Frequency of maximum amplitude (Hz) | frequency | 1.37 | <0.001 | 1.20 | <0.05 |
| Unit 1 Q25 (Hz) | frequency | 1.14 | <0.001 | 1.06 | NS |
| Unit 1 Q50 (Hz) | frequency | 1.25 | <0.001 | 0.94 | NS |
| Unit 1 Q75 (Hz) | frequency | 1.22 | <0.001 | 0.93 | NS |
| Unit 1 E1200 Hz (%) | frequency | 1.24 | <0.001 | 1.05 | NS |
| Down-movement part |  |  |  |  |  |
| Duration (s) | temporal | 1.16 | <0.001 | 0.94 | NS |
| No. of units | temporal | 1.42 | <0.001 | 1.07 | NS |
| Fundamental frequency (Hz) | frequency | 1.89 | <0.001 | 1.17 | <0.05 |
| Frequency of maximum amplitude (Hz) | frequency | 1.07 | <0.001 | 1.94 | <0.01 |
| Q25 (Hz) | frequency | 1.27 | <0.001 | 1.47 | <0.01 |
| Q50 (Hz) | frequency | 1.52 | <0.001 | 1.68 | <0.01 |
| Q75 (Hz) | frequency | 1.51 | <0.001 | 1.32 | <0.05 |
| Energy below 1200 Hz (%) | frequency | 1.45 | <0.001 | 1.65 | <0.01 |
| Unit 1 Duration (s) | temporal | 2.02 | <0.001 | 1.37 | <0.05 |
| Unit 1 No. of segments | temporal | 1.51 | <0.001 | 1.70 | NS |
| Unit 1 Pulse Rate (Hz) | temporal | 1.23 | <0.001 | 1.29 | NS |
| Unit 1 Fundamental Frequency (Hz) | frequency | 1.51 | <0.001 | 1.19 | NS |
| Unit 1 Frequency of maximum amplitude (Hz) | frequency | 1.07 | <0.001 | 1.11 | <0.05 |
| Unit 1 Q25 (Hz) | frequency | 1.09 | <0.001 | 1.25 | NS |
| Unit 1 Q50 (Hz) | frequency | 1.31 | <0.001 | 1.25 | <0.05 |
| Unit 1 Q75 (Hz) | frequency | 1.26 | <0.001 | 1.30 | <0.05 |
| Unit 1 E1200 Hz ( %) | frequency | 1.27 | <0.001 | 1.19 | <0.05 |

Since a call is a single unit from which we measured different variables, the potential individual signatures were more realistic when considering a combination of acoustic variables. The RF classification for individuals showed a global accuracy of prediction of 90% for females and 76% for males. The percentage of correct classification varied between 40% and 100% depending on individuals, with a median value of 80% for males and 86% for females (Table 2). These predictions were between 5.9 and 14.3 times better than a prediction by chance.

Interestingly, the most important variable to discriminate individuals was different depending on the sex of the gannets (Fig. 4). In both cases, the IOIm (units’ temporal rate) together with a frequency variable were the two most important variables to distinguish individuals. For females, the Fmax in the down-movement part was important, followed by the IOIm. For males, the IOIm was the most important, followed by the F0 in the down-movement part.

**Figure 4.**
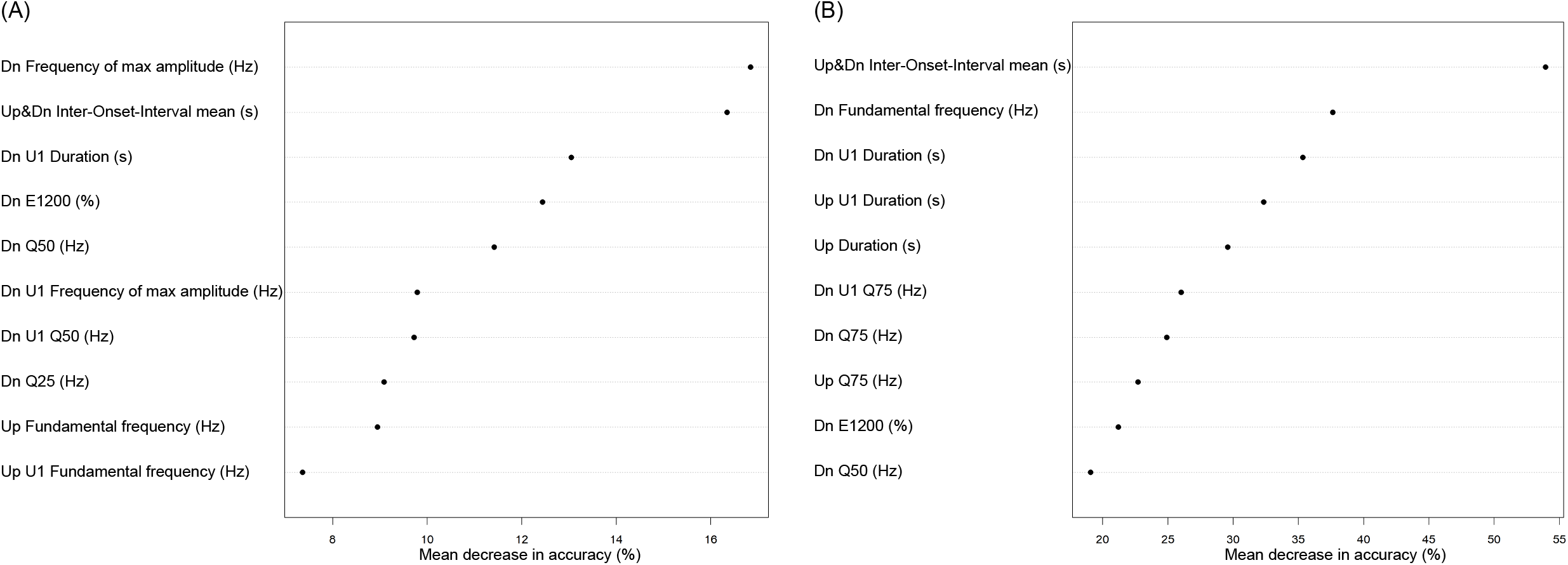
Analyses on individual vocal signatures among females (A) and among males (B) in the display calls of Cape gannets. Ranking of the importance of acoustic variables for individual distinction calculated as a mean decrease in accuracy in the random forest algorithm. Only the first 10 variables are shown here. Up = up-movement part. Dn = downmovement part. U1 = first unit. Q25, Q50, Q75 = first, second and third quartile of energy distribution. E1200 = percentage of energy occurring below 1200 Hz.

## 5. Discussion

This paper presents an exhaustive description of the acoustic structure of the display call in adult, nesting Cape gannets. We showed that the vocalisations associated with the characteristic up and down head movement behaviour could be used reliably for identification of individuals and sex. Both the frequency variables (mostly fundamental frequency, but also frequency of maximum amplitude) and a measure of the temporal rate of unit production within a vocalisation (IOI Inter-Onset-Interval, [42]) were the most important variables to discriminate sexes and individuals. Furthermore, our findings clearly demonstrate that the sex of Cape gannets can be identified directly in the field using non-invasive methodology, as opposed to retrospective costly and timely genetic analyses.

### Sexual dimorphism in display calls

Three quarters of the total number of calls recorded from molecular-sexed gannets were from males (74 out of 97). Since the display calls are associated with a territorial behaviour [31], this suggests that males express a more territorial behaviour at their nests than females. In seabirds, such differences between females and males are common and could potentially explain the greater occurrence of male vocalisations [49,50].

The display calls produced by female and male Cape gannets can be differentiated by a combination of both temporal and spectral acoustic variables, thus the sex information is based on a multi-parametric coding of the call. More specifically, we found that two acoustic variables were clearly discriminating between the two sexes in Cape gannets: the fundamental frequency and the temporal Inter-Onset-Interval between successive sound units within a call. The difference observed in fundamental frequency, with females displaying a lower fundamental frequency value than males, could result from differences in the anatomy of the vocal apparatus [51,52] and/or differences in sexual hormones [53]. In addition, female calls on average consisted of more units, even though the total duration of their calls did not vary significantly from males, which demonstrates that females call at a faster rate (Supplementary Audios 1-4). The potential drivers of these differences in call rate, however, remain unclear.

In gannet species, anecdotal evidence for differences in the vocalisations between sexes in Cape gannets and northern gannets has been suggested before [54] but has never been thoroughly investigated in either of the two species. In Australasian gannets, sexual differences were not found in a variety of different call elements [30]. However, the authors did not measure any temporal parameters or the fundamental frequency, limiting the ability for comparison with our study.

Vocal dimorphism has been found in a number of seabird species, where some acoustic parameters substantially vary between the different sexes. Differences in the fundamental frequency between sexes have been found in other species such as black-legged kittiwakes [27], yelkouan shearwaters [29] and king penguins [19]. Differences in the temporal rate of sound production between sexes has not been commonly studied in seabirds (but see [55]) but other temporal parameters including the duration of different parts of the call (sound units or silences between successive units) have been shown to be sexually dimorphic in blacklegged kittiwakes [27] and yelkouan shearwaters [8].

The observed differences in the calls between males and females could be used to determine the sex of an individual in the field, therefore using a method that is non-invasive, more efficient and less costly compared to currently-used genetic analyses. Auditory recognition (based on the temporal rate of units) would require some training but seems feasible, as has been shown for petrels [56], prions [57], shearwaters [58] and penguins [19]. Alternatively, reliable sexing can certainly be achieved through recording vocalisations in the field and using signal processing software to measure the two discriminating variables (IOI and fundamental frequency). The use of both variables simultaneously seems necessary to avoid the potential overlapping values between males and females (Fig. 3). In addition, the recording of a few vocalisations (e.g. 2-3 calls per individual) is probably necessary to further reduce potential confusion and errors. Indeed, we also observed intra-individual variations in the vocalisations, so that if the measures on a particular call may unfortunately fall within the overlap area, the repetitive measures of several calls will most probably allow for a reliable sex-identification.

### Individual signatures in display calls

This study provided quantitative evidence of individual signatures in the display calls of adult breeding Cape gannets, which most likely plays an important role in individual recognition in these large breeding colonies [35,37]. Two acoustic features appeared to contribute the most to differentiate individuals, the Inter-Onset-Interval (related to the temporal rate of unit production) and frequency parameters during the down-movement part (Fmax for females and F0 for males). Spectral differences, such as the fundamental frequency and energy distribution in seabird vocalisations have been associated with anatomical differences in their airways [59,60] according to the source-filter theory [61]. Slight differences in vocal tract anatomy between individuals most likely explain the differences in the fundamental frequency between individuals we found in this study. It remains unclear if the differences in the temporal rate could also result from differences in the anatomy among the different individuals, or if it could be related to differences in body condition, hormone levels, motivation or personality [53].

In northern gannets, differences between individuals were evident in the envelopes of their landing calls [35]. Individual signatures have also been found in the frequency parameters of the nesting vocalisations of Australasian gannets [30]. The results of these studies [30,35,this study] demonstrate that individual recognition might be essential in breeding colonial gannets, and that this recognition could be largely based on acoustic signals.

Individual signatures are common in the vocalisations of nesting colonial seabirds [17,27,60], although the signatures can be carried on a variety of acoustic variables. Individual vocal signatures were found in the greeting calls of black-legged kittiwakes, on both temporal and frequency features [27]. In the yelkouan shearwaters, individual signatures were identified in the display calls and were particularly evident when looking at temporal parameters [8]. In blue-footed boobies, individual discrimination was sufficient using only spectral features for females, however individual discrimination in males required both temporal and spectral features [18]. Temporal parameters seem to be important for individual vocal signatures in other seabird species, emphasising the potential importance in individual signatures in gannet species.

## 6. Conclusion

Cape gannets breed in large and dense colonies and most likely rely on a combination of signals to identify individuals as well as the opposite sex. This study demonstrated that sexual and individual signatures are carried in their display call, and potentially provides a valuable tool for identification in the field, which is important for population monitoring and conservation. The temporal rate of unit production within a display call played a primary role for both sexual and individual discrimination, suggesting this aspect should be considered more often in non-passerines species.

## Supporting information

Supplementary Video 1. Display call of a male Cape gannet (individual 17U)

Supplementary Audio 1. Display call of a male Cape gannet (individual 17U)

Supplementary Audio 2. Display call of a female Cape gannet (individual 19U)

Supplementary Audio 3. Display call of a male Cape gannet (individual 15L)

Supplementary Audio 4. Display call of a female Cape gannet (individual 15R)

## 7. Acknowledgements

We thank South African National Parks for logistical support during fieldwork. We thank Rabi’a Ryklief, Jonathan Botha and Melanie Wells for their help in the field.

## 8. Ethical Statement

Research using animals must adhere to local guidelines and state that appropriate ethical approval and licences were obtained.

## 11. Authors’ Contributions

KBF: investigation, formal analysis, writing - original draft, project administration; AT: conceptualization, methodology, investigation, supervision, data curation, formal analysis, writing – original draft; MC: formal analysis, methodology, writing – review & editing; TA: conceptualization, methodology, resources, writing – review & editing; IC: conceptualization, methodology, resources, writing – review & editing; PAP: supervision, resources, writing – review & editing, funding acquisition.

## 7. Supporting Information

Supplementary Figure 1. Convergence of the random forest algorithm observed as a decrease and stabilisation of global error of prediction as a function of the number of trees grown for the three random forest analyses: (A) comparing calls between sexes (females in red, males in green and global error in black), (B) comparing calls among female individuals (each individual represented by a different colour, with the global error in black), (C) comparing calls among male individuals (each individual represented by a different colour, with the global error in black).

Supplementary Video 1. Display call of a male Cape gannet (individual 17U).

Supplementary Audio 1. Display call of a male Cape gannet (individual 17U). Supplementary Audio 2. Display call of a female Cape gannet (individual 19U).

Supplementary Audio 3. Display call of a male Cape gannet (individual 15L). Supplementary Audio 4. Display call of a female Cape gannet (individual 15R).

